# ARID1A loss derepresses human endogenous retrovirus-H to modulate BRD4-dependent transcription

**DOI:** 10.1101/2021.06.28.450127

**Authors:** Chunhong Yu, Xiaoyun Lei, Fang Chen, Song Mao, Lu Lv, Honglu Liu, Xueying Hu, Runhan Wang, Licong Shen, Na Zhang, Yang Meng, Yunfan Shen, Pishun Li, Shi Huang, Hao Shao, Changwei Lin, Zhuohua Zhang, Kai Yuan

**Affiliations:** Hunan Key Laboratory of Molecular Precision Medicine, Department of Oncology, Xiangya Hospital, Central South University, Changsha, Hunan, China; Hunan Key Laboratory of Medical Genetics, School of Life Sciences, Central South University, Changsha, Hunan, China; Department of Gynecology, Xiangya Hospital, Central South University, Changsha, Hunan, China; National Clinical Research Center for Geriatric Disorders, Xiangya Hospital, Central South University, Changsha, Hunan, China; Department of Gastrointestinal Surgery, The Third Xiangya Hospital, Central South University, Changsha, Hunan, China; The Biobank of Xiangya Hospital, Central South University, Changsha, Hunan, China

## Abstract

The transposable elements (TEs) through evolutionary exaptation have become an integral part of human genome, offering ample regulatory sequences and shaping chromatin 3D architecture. While the functional impacts of TE-derived sequences on early embryogenesis are recognized, their role in malignancy has only started to emerge. Here we show that many TEs, especially the pluripotency-related endogenous retrovirus H (HERVH), are abnormally activated in colorectal cancer (CRC) samples. The transcriptional upregulation of HERVH is associated with mutations of several tumor suppressors including ARID1A. Knockout of ARID1A in CRC cells leads to increased accessibility at HERVH loci and enhanced transcription, which is dependent on ARID1B. Suppression of HERVH in CRC cells and patient-derived organoids impairs tumor growth. Mechanistically, HERVH transcripts colocalize with nuclear BRD4 foci, modulate their dynamics, and co-regulate many target genes. Altogether, we uncover a critical role for ARID1A in restraining HERVH, which can promote tumorigenesis by stimulating BRD4-dependent transcription when ARID1A is mutated.

## Introduction

We have been facing constant viral attacks during the course of evolution. While most viruses come and go, few have invaded and colonized the germline genome, becoming a significant fraction of transposable elements (TEs) that contribute more than 50% to the human nuclear DNA content^1–4^. Human TEs include DNA transposons, long terminal repeat (LTR) retrotransposons, and non-LTR retrotransposons. The majority of them has lost the ability to transpose during evolution and had long been regarded as functionless repetitive DNA. Recent studies however have begun to reveal that TEs are an abundant source of many regulatory sequences^2, 3, 5, 6^, such as microRNAs (miRNAs) and long noncoding RNAs (lncRNAs)^7–12^, and that TEs are co-opted to serve important functions including transcriptional regulation, chromatin organization and 3D compartmentalization, especially during early embryogenesis and in embryonic stem cells (ESCs)^6, 13–23^.

The endogenous retroviruses (ERVs), which have been identified half a century ago^24^, make up 8% of the human genome. They are LTR retrotransposons and have similar compositions to retroviruses, with internal coding sequences (gag-pro-pol-env) flanked by a pair of identical LTRs containing cis-regulatory elements for transcription. By estimation, there are 98,000 copies of ERVs and their derivatives, with human endogenous retrovirus H (HERVH) being one of the most abundant groups, comprising a total of ∼2000 copies among which ∼100 are close to full-length^25, 26^. ERVs are largely in heterochromatin and transcriptionally repressed by an expanding battery of epigenetic mechanisms^4, 27, 28^, including methylation of histone H3 on lysine 9 (H3K9) or lysine 27 (H3K27), DNA methylation, as well as the RNA N(6)-methyladenosine (m(6)A) modification^29, 30^. Of note, these regulatory mechanisms are often redundant and function in a context-specific manner^31, 32^, reflecting the sophisticated evolutionary arms race between viral sequences and the host genome^2–4^.

ERVs are not always inactive. During the profound epigenetic resetting in early embryonic development, ERVs are systematically transcribed in a stage-specific manner, coinciding with different cellular identities and differentiation potencies^33^. While a comprehensive understanding of ERVs function during early embryogenesis is yet to be established, recent studies have revealed the intimate relationship between HERVH and the human pluripotency network^19^. Depending on different variants of LTR (LTR7, LTR7Y, and LTR7A/B/C), the transcription of HERVH internal sequence (HERVH-int) is activated from 4-cell stage to blastocyst^33^. HERVH transcripts are also highly abundant in human ESCs as well as induced pluripotent stem cells (iPSCs), and moreover, the naïve-like pluripotency is associated with higher levels of HERVH expression^15, 16, 19, 26^. Activation of HERVH promotes both the acquisition and maintenance of pluripotent states, by generating noncoding RNAs (ncRNAs) or producing chimeric transcripts with protein-coding genes via alternative splicing^15, 16^. The transcriptionally active HERVH can also demarcate topologically associated domains (TADs) and help maintain a pluripotent chromatin architecture^22^. Cancer development in many aspects parallels the process of early embryogenesis. This includes regain the capacity of self-renewal and dramatic alterations in epigenetic landscapes. Interestingly, reactivation of HERVH is also observed in several types of human cancer, such as colorectal carcinomas (CRCs)^34–39^, however, a mechanistic insight into this reactivation is lacking and its functional consequence unclear.

The SWI/SNF (mating type SWItch/Sucrose NonFermentable) family chromatin remodelers, BAF, PBAF, and GBAF, regulate chromatin packing and transcription by controlling the dynamics of nucleosomes^40^. As a subunit of the BAF complex, ARID1A functions as a bona fide tumor suppressor and is mutated in approximately 8% of all human cancers^40–44^. Mutation of ARID1A sensitizes cancer cells to bromodomain and extraterminal domain (BET) inhibitors^45, 46^, likely due to its indispensable role in maintaining normal enhancer function by influencing BRD4 activity^42, 46, 47^. How ARID1A mutation affects BRD4 remains unknown. Here, we show that loss of ARID1A results in an ARID1B-dependent upregulation of HERVH, whose transcripts partition into nuclear BRD4 foci and contribute to the BRD4-dependent gene regulatory network. This ARID1B-HERVH-BRD4 axis is crucial for the growth of CRC cells and patient-derived organoid, offering novel treatment opportunities for ARID1A mutated cancers.

## Results

### HERVH is abnormally upregulated in CRCs

The majority of the human genome is comprised of various repetitive DNA sequences, most of which are transcribable. To globally characterize the expression of repetitive DNA elements in CRCs, we collected 521 colon adenocarcinoma (COAD) and 177 rectum adenocarcinoma (READ) RNA-seq data from The Cancer Genome Atlas (TCGA), filtered and grouped them according to the variables (Fig. S1A), and quantified the repeats expression using the human RepeatMasker Repeats annotation (https://genome.ucsc.edu/cgi-bin/hgTables). We first applied principal component analysis (PCA) to the gene expression as well as the repeats expression data from 51 normal and 631 tumor samples (Fig. S1A). Both the genes and repeats showed distinct expression profiles that successfully demarcated the normal and tumor samples (Fig. 1A-1B and Supplementary Table 1). We next categorized the differentially expressed repeats. While the simple repeats were the most abundant, many LTR retrotransposons (also known as ERVs) showed altered expression between normal and tumor tissues (Fig. 1C and Supplementary Table 1). Of the 580 ERVs, 44 were downregulated and 84 upregulated in CRC tumor tissues (Fig. 1D and Supplementary Table 1). To validate these upregulated ERVs, we repeated the analysis with another independent RNA-seq dataset (GSE50760) of CRC tissues, and identified 17 ERVs that showed consistent upregulation. Specific activation of ERVs is linked with pluoripotency in embryonic cells^6, 20^. We compared the upregulated ERVs in CRC tissues with that observed in early embryos and ESCs, and pinpointed two elements, HERVH-int and LTR7Y, that constitute a full-legth HERVH (Fig. 1E and Supplementary Table 1). Both elements showed increased expression in tumor tissues and their expressions were highly correlated with each other in the TCGA colorectal dataset (COREAD) (Fig. S1B-S1D and Supplementary Table 1). To further confirm the upregualtion of HERVH in CRCs, we performed RNAscope analysis for HERVH RNA on CRC tissue array. Compared with the matched peritumoral tissues, the tumoral tissues showed significantly stronger RNAscope signals (Fig. S1E-S1F). We then investigated the association of the expression levels of HERVH with the clinical outcomes of the CRC patients using the TCGA COREAD dataset, and observed that higher HERVH-int expression predicted poorer survival (Fig. 1F and Supplementary Table 1).

**Figure 1.**
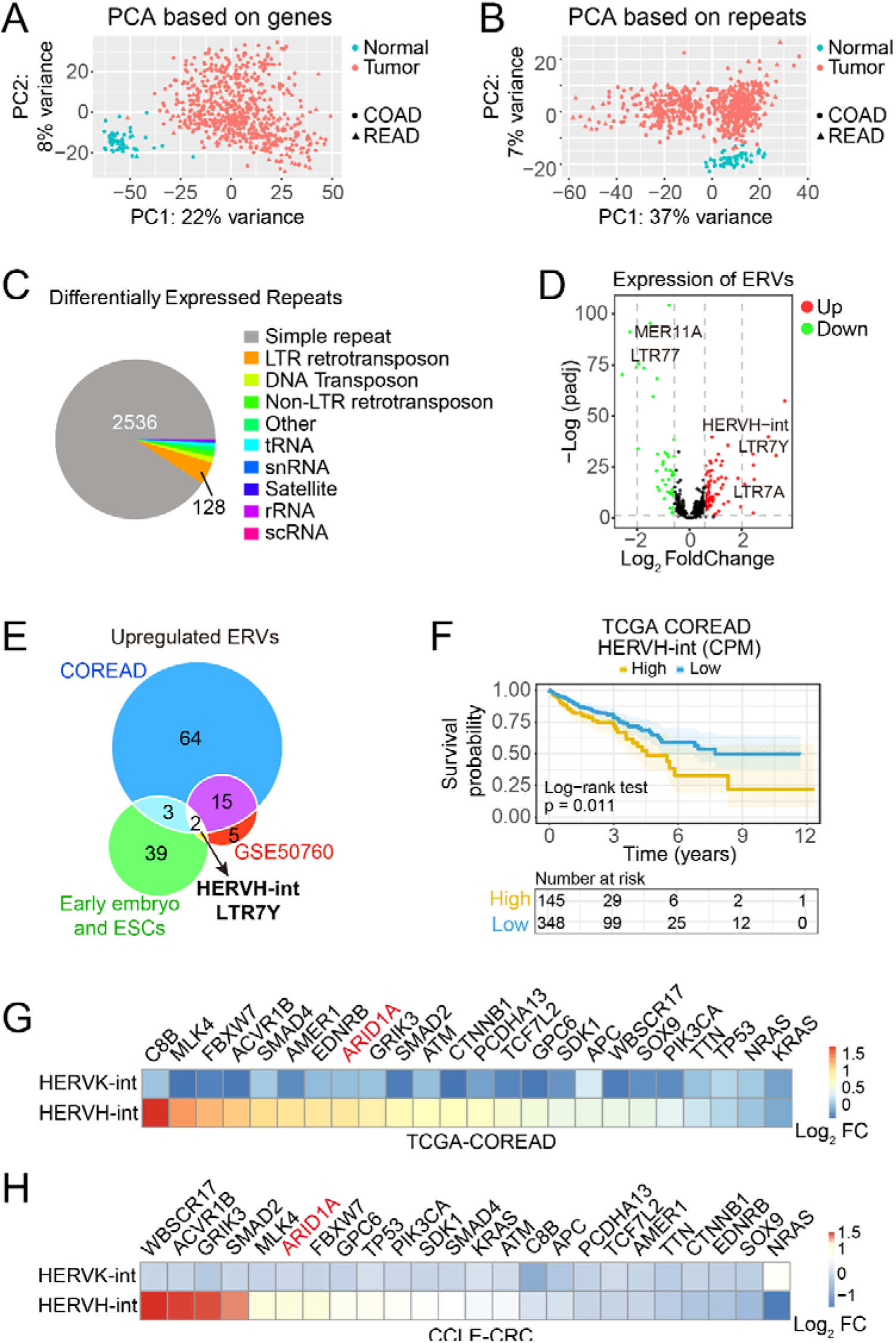
Characterization of HERVH expression in CRCs. (A) Principal component analysis (PCA) of gene expression of 51 normal and 631 CRC tumor tissues from the TCGA COREAD dataset. (B) PCA based on the expression of repetitive sequences in the same TCGA dataset. (C) Classification of differentially expressed repetitive sequences (adjusted p-value < 0.05 and |Log_2_ FoldChange| > 0.585). (D) Volcano plot of differentially expressed ERVs. Up (red) and down (green) regulated ERVs are determined with the cut-off values of adjusted p-value < 0.05 and |Log_2_ FoldChange| > 0.585. (E) Overlap analysis of upregulated ERVs in CRCs samples and early embryonic cells identifies the internal coding sequences of HERVH (HERVH-int) and its corresponding LTR (LTR7Y) as the commonly upregulated elements. (F) Survival analysis based on the expression level of HERVH-int and the overall survival (OS) from 493 patients with AJCC pathologic tumor stage greater than I. The mean expression value of HERVH-int is used to demarcate the HERVH-int-High (145 patients) and HERVH-int-Low (348 patients) groups. (G) Correlation analysis of HERVH-int expression and mutational status of the most frequently mutated genes in CRCs using the TCGA dataset. (H) Correlation of HERVH-int expression and gene mutations in CRC cell lines from the CCLE dataset.

**Figure S1.**
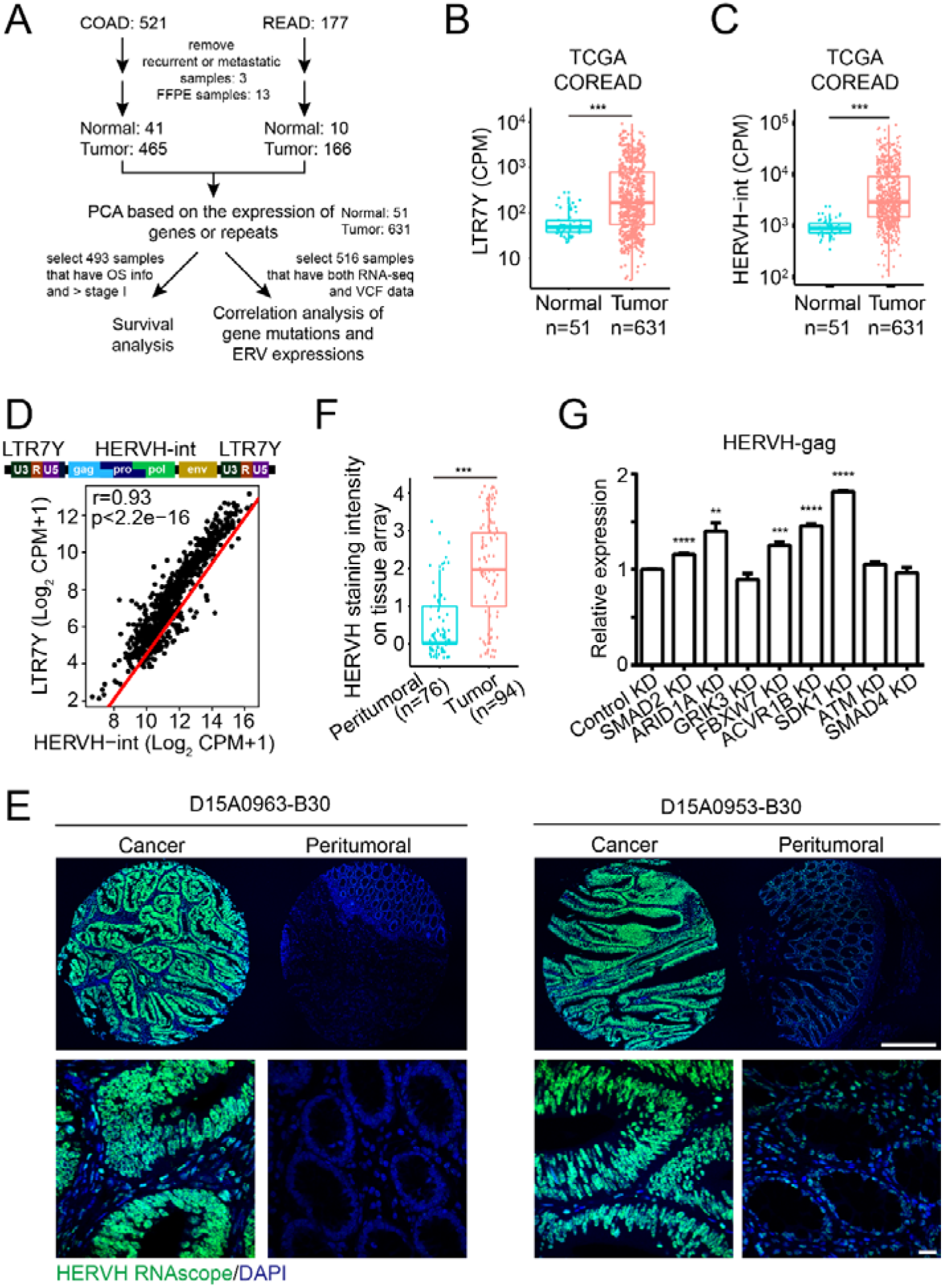
Expression of HERVH in CRC samples. (A) The inclusion and exclusion criteria for the TCGA-COREAD samples used in Figure 1. (B-C) Box plots of the expression of LTR7Y and HERVH-int in the TCGA-COREAD dataset. ***p < 0.001 by Wilcox test. (D) Correlation of the expression of HERVH-int and LTR7Y in the TCGA-COREAD dataset. The Pearson correlation coefficient (r) and the p-value are shown. (E) Representative images of RNAscope staining of HERVH transcripts on CRC tissue array. Bars: 500 μm in the upper panels and 20 μm in lower insets. (F) Quantification of the RNAscope signals from the peritumoral and tumor tissues on the CRC tissue array. ***p < 0.001 by Wilcox test. (G) qPCR analysis of HERVH expression in cells treated with the indicated siRNA. **p < 0.01, ***p < 0.001, ****p < 0.0001 by t-test.

Molecular characterization of CRC samples have revealed 24 genes that are significantly mutated^48^. To interrogate the relationship between these gene mutations and the upregulation of HERVH, we selected 516 CRC samples with genetic variation data from the TCGA COREAD dataset, extracted their mutational signatures, and correlated the mutational status of one of the 24 genes with the expression of either HERVH-int or HERVK-int for comparison (Fig. S1A). In contrast to HERVK-int whose expression showed no obvious association with any gene mutations analyzed, the expression of HERVH-int correlated with the mutational status of several genes (Fig. 1G and Supplementary Table 1). We expanded this analysis to the 59 CRC cell lines in cancer cell line encyclopedia (CCLE)^49^ (Fig. 1H and Supplementary Table 1), and identified a list of genes whose mutation was consistently correlated with upregulation of HERVH. This included MLK4, FBXW7, ACVR1B, ARID1A, GRIK3, and SMAD2.

### Loss of ARID1A leads to transcriptional activation of HERVH

Knockdown the expression of some of the listed genes by small interfering RNA (siRNA) resulted in increased transcription of HERVH (Fig. S1G). We selected ARID1A for further functional validation, because it is a DNA-binding subunit of the BAF chromosome remodeler complex and its inactivation mutations occur in a broad spectrum of human cancers^40, 41, 43^.

To comprehensively depict the changes of repeats expression upon ARID1A loss, we collected and analyzed two independent RNA-seq data of HCT116 wild type (WT) and its isogenic ARID1A knockout (KO) cell lines^42, 50^. Of note, the LTR retrotransposons or ERVs were the most upregulated repeat group in ARID1A KO cells (Fig. 2A, S2A, and Supplementary Table 2). We ranked all the ERVs according to their fold changes (Fig. 2B, S2A, and Supplementary Table 2). HERVH-int and two of its associated LTRs, LTR7Y and LTR7, were the three most significantly upregulated elements, whereas other HERVH-related LTRs didn’t show consistent upregulation (Fig. 2C, S2A, and Supplementary Table 2). Scatter plots of the expression of all 580 ERVs revealed that HERVH-int, LTR7Y, and LTR7 were already expressed in HCT116 WT cells but the ARID1A inactivation further increased their abundance (Fig. 2D, S2C, and Supplementary Table 2). These observations were further validated with our own RNA-seq data of ARID1A WT and KO HCT116 cells (Fig. S2B, S2D, and Supplementary Table 2). Overlapping the significantly upregulated ERVs in the three datasets spotted HERVH-int and its LTR as the only unambiguously activated elements upon ARID1A loss (Fig. 2E). We generated additional ARID1A KO colorectal cell lines to further confirm the observed upregulation of HERVH (Fig. 2F). qPCR analyses with primers specifically targeting the *gag* and *pol* sequences of HERVH-int revealed increased transcripts abundance in all three ARID1A KO cell lines (Fig. 2G and 2H). To test if the transcriptional activation of HERVH can be suppressed by re-expression of ARID1A, we infected the ARID1A KO cells with lentiviruses carrying ARID1A or its counterpart ARID1B^51, 52^ (Fig. 2I-2K). The re-introduction of ARID1A significantly downregulated the expression of HERVH. Interestingly, overexpression of ARID1B didn’t rescue but instead mildly increased the amount of HERVH transcripts in ARID1A KO cells (Fig. 2K).

**Figure 2.**
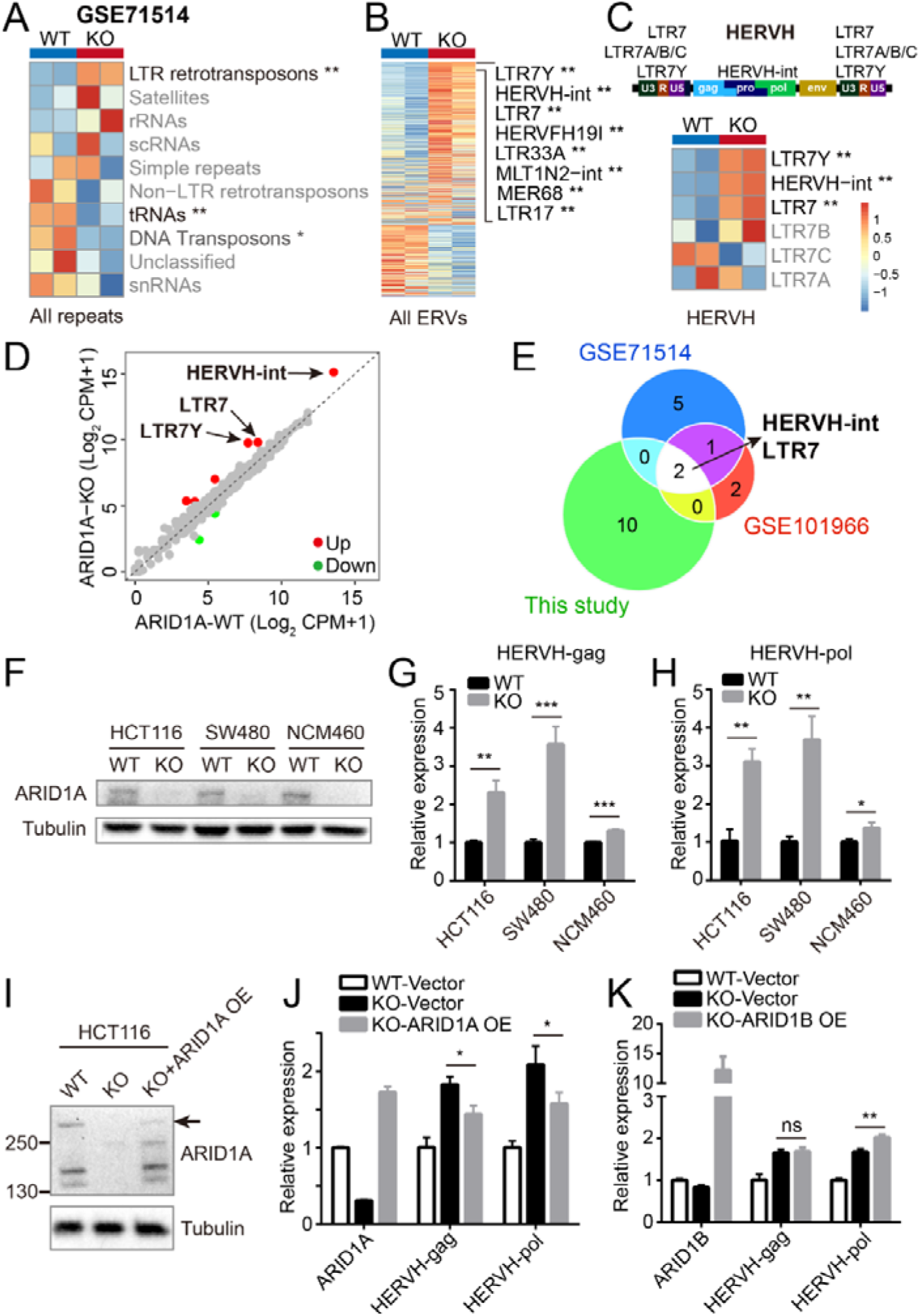
ARID1A loss leads to upregulation of HERVH. (A-C) Heatmaps of the expression of different repetitive sequences in wild type (WT) and ARID1A knockout (KO) HCT116 cells. The differential expression is tested based on a model using the negative binomial distribution (adjusted p values are labeled as *padj < 0.05, **padj < 0.01). The schematic of a typical full-length HERVH element is shown in (C). (D) A scatter plot of the expression of all 580 ERVs in WT and KO HCT116 cells. The up- or downregulated ERVs are labeled in red or green respectively (|Log_2_ FoldChange| > 1 and adjusted p-value < 0.05). (E) Venn diagram showing that HERVH is repetitively upregulated in three independent sequencing experiments with HCT116 ARID1A WT and KO cells. (F) Western blots of different ARID1A WT and KO cell lines. (G-H) qPCR analyses of HERVH expression in ARID1A WT and KO cell lines using two different primer sets. *p < 0.05, **p < 0.01, ***p < 0.001 by t-test. (I) Western blots showing ARID1A protein levels in WT, KO, and ARID1A rescued KO HCT116 cells. Arrow indicates the full-length ARID1A band. (J) qPCR analysis of ARID1A and HERVH expression in the indicated groups of cells. *p < 0.05 by t-test. (K) qPCR analysis of ARID1B and HERVH expression. **p < 0.01 by t-test.

**Figure S2.**
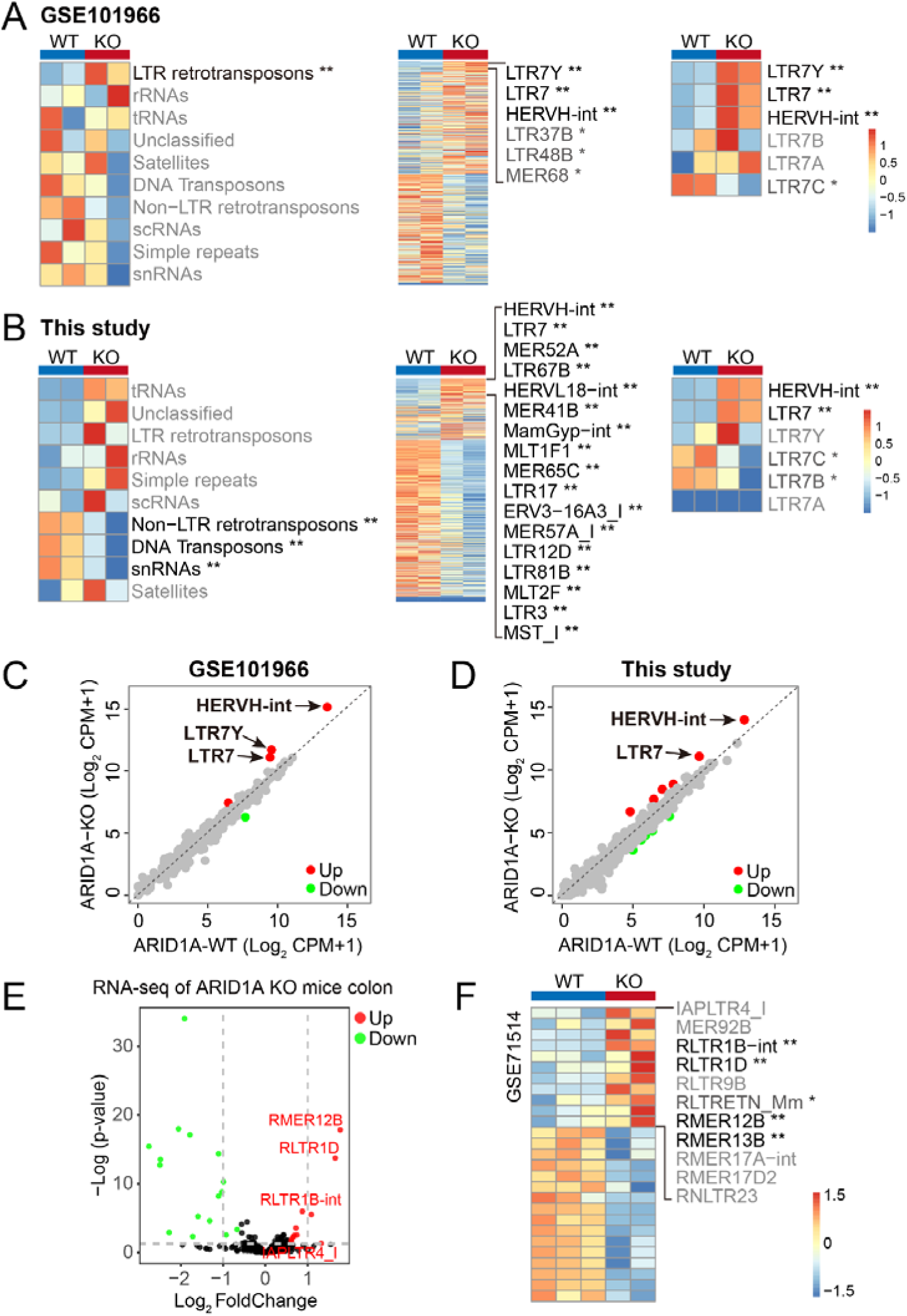
ARID1A loss derepresses ERVs. (A-B) Heatmaps of the expression of different repetitive sequences in wild type (WT) and ARID1A knockout (KO) HCT116 cells using our own RNA-seq data and that from GSE101966. *padj < 0.05, **padj < 0.01. (C-D) Scatter plots of the expression of all 580 ERVs in WT and ARID1A KO HCT116 cells using different datasets. (E) Volcano plot of differentially expressed ERVs in WT and ARID1A KO mouse colons. Up (red) and down (green) regulated ERVs are determined with the cut-off values of adjusted p-value < 0.05 and |Log_2_ FoldChange| > 0.585. (F) Heatmap showing differentially expressed mouse ERVs in ARID1A KO mouse colon using GSE71514 dataset. *padj < 0.05, **padj < 0.01.

Unlike genes, ERVs are quite diverse between primates and rodents. To assess the effect of ARID1A inactivation on ERVs expression in mice, we analyzed RNA-seq data of the colon epithelial cells from WT or ARID1A KO mice^42^. Of the 423 ERVs in the mouse genome, 16 were downregulated and 11 upregulated in the absence of ARID1A (Fig. S2E and Supplementary Table 3). The upregulated ERVs included RLTR1B-int, RLTR1D, RMER12B, IAPLTR4_I (Fig. S2F and Supplementary Table 3). Therefore, the influence of ARID1A on ERVs seemed to be universal, and in human, the most responsive element toward ARID1A mutation was the HERVH.

### Transcription of HERVH in the absence of ARID1A is dependent on ARID1B

To investigate the mechanism of how ARID1A loss induced HERVH transcription, we put our focus on ARID1B, which shares 60% homology with ARID1A^51^. Functioning as the rigid structural core, they are mutually exclusive in the BAF complex^52, 53^. ARID1B is essential for the survival of ARID1A mutated cancer cells, by supplying residual BAF complex activities to maintain chromatin accessibility at enhancers and regulate RNA polymerase II dynamics^50, 54, 55^.

We first examined the influence of ARID1B on the expression of repetitive elements using published RNA-seq data^50^ (Fig. 3A and Supplementary Table 4). Knocking down the expression of ARID1B by short hairpin RNA (shRNA) in WT HCT116 cells (WT-KD) only showed limited effects on repeats expression, however, ARID1B knockdown in ARID1A KO cells (KO-KD) dramatically reduced the transcripts abundance of many repeats, especially the LTR retrotransposons (ERVs) (Fig. 3B and Supplementary Table 4). Of note, the upregulation of several HERVH elements was partially reversed by ARID1B KD (Fig. 3C and Supplementary Table 4). Using two different shRNAs targeting ARID1B, we verified the suppression of HERVH by ARID1B KD in ARID1A KO cells (Fig. 3D). Both ARID1A and ARID1B harbor DNA-binding activity^51^. We next analyzed their occupancy on the HERVH elements using ChIP-qPCR (Fig. S3A and S3B). While the amount of ARID1A on HERVH was minimized in ARID1A KO cells, the binding of ARID1B to HERVH was compensatorily increased, maintaining a comparable amount of BAF activity at HERVH loci in these cells (Fig. S3C). The ARID1A- and ARID1B-containing BAF complexes are associated with different histone acetyltransferase (HAT) and deacetylase (HDAC) activities^52^. To characterize the epigenetic changes accompanying this subunit switch of BAF complex on HERVH, we identified the commonly derepressed genomic HERVH loci in different datasets and analyzed their chromosome accessibility as well as histone modifications (Fig. S3D and Supplementary Table 5). In ARID1A KO HCT116 cells, we observed some increase in accessibility at the HERVH loci (Fig. 3E, 3H, and Supplementary Table 5). Interestingly, acetylation of H3K27 (H3K27ac), as well as mono methylation of histone 3 on lysine 4 (H3K4me), was also increased (Fig. 3F-3H, and Supplementary Table 5). We further validated this increase of H3K27ac on HERVH by ChIP-qPCR (Fig. S3E). To test whether HDACs and HATs contributed to the dysregulation of HERVH in the absence of ARID1A, we treated WT HCT116 cells with two different HDAC inhibitors, SAHA and TSA. Both inhibitors stimulated the expression of HERVH (Fig. S3F). We treated the ARID1A KO cells with MG149, an inhibitor of Tip60 which is a HAT associated with ARID1B-BAF complex^52^. The inhibition of Tip60 suppressed the activation of HERVH upon ARID1A loss (Fig. S3G). To identify which transcription factor (TF) accounted for the increased expression of HERVH, we examined the expression levels of all the TFs that are able to bind HERVH in ARID1A KO HCT116 cells (Fig. S3H), individually knocked down their expression by siRNA, and performed RNA-seq analysis (Fig. S3I and Supplementary Table 6). Only SP1 knockdown significantly reduced the expression of HERVH (Fig. S3J).

**Figure 3.**
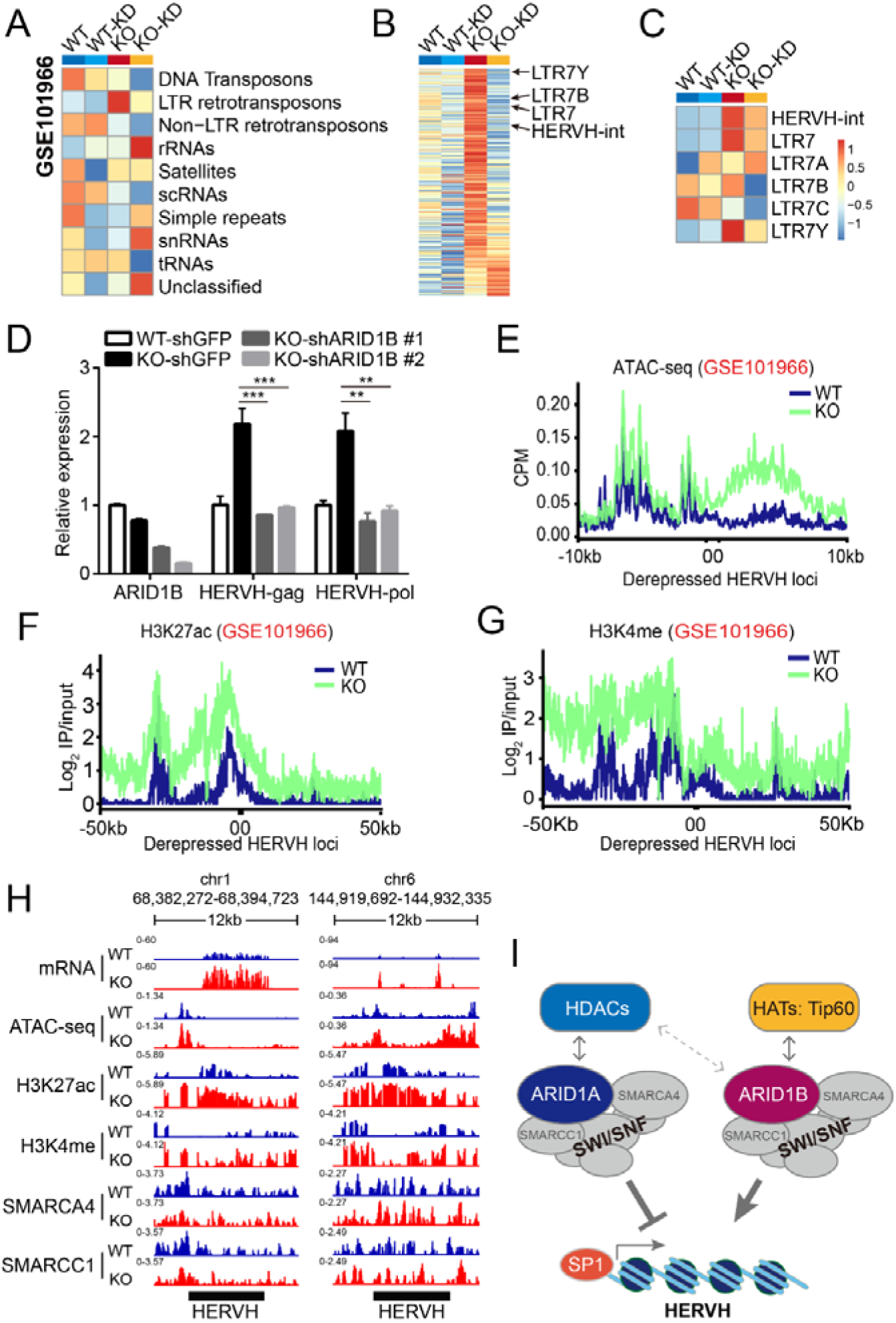
ARID1B is required for the upregulation of HERVH in the absence of ARID1A. (A-C) Heatmaps of the expression of different repetitive sequences in WT or ARID1A KO HCT116 cells treated with scrambled control or ARID1B (-KD) shRNAs. (D) qPCR results showing the expression levels of ARID1B and HERVH in ARID1A WT and KO cells treated with shRNA targeting GFP or ARID1B. **p < 0.01, ***p < 0.001 by t-test. (E) ATAC-seq results from ARID1A WT and KO HCT116 cells (GSE101966) demonstrate increased chromatin accessibility at the derepressed HERVH loci in ARID1A KO cells. (F-G) ChIP-seq data (GSE101966) show increased H3K27ac and H3K4me at the derepressed HERVH loci in ARID1A KO HCT116 cells. (H) Genomic snapshots of RNA-seq, ATAC-seq, and ChIP-seq signals at two representative HERVH loci. (I) Model showing the mutually exclusive relationship between ARID1A- and ARID1B-containing BAF complexes, their differential associations with HDACs and HATs, and the different regulatory functions imposed on HERVH loci.

**Figure S3.**
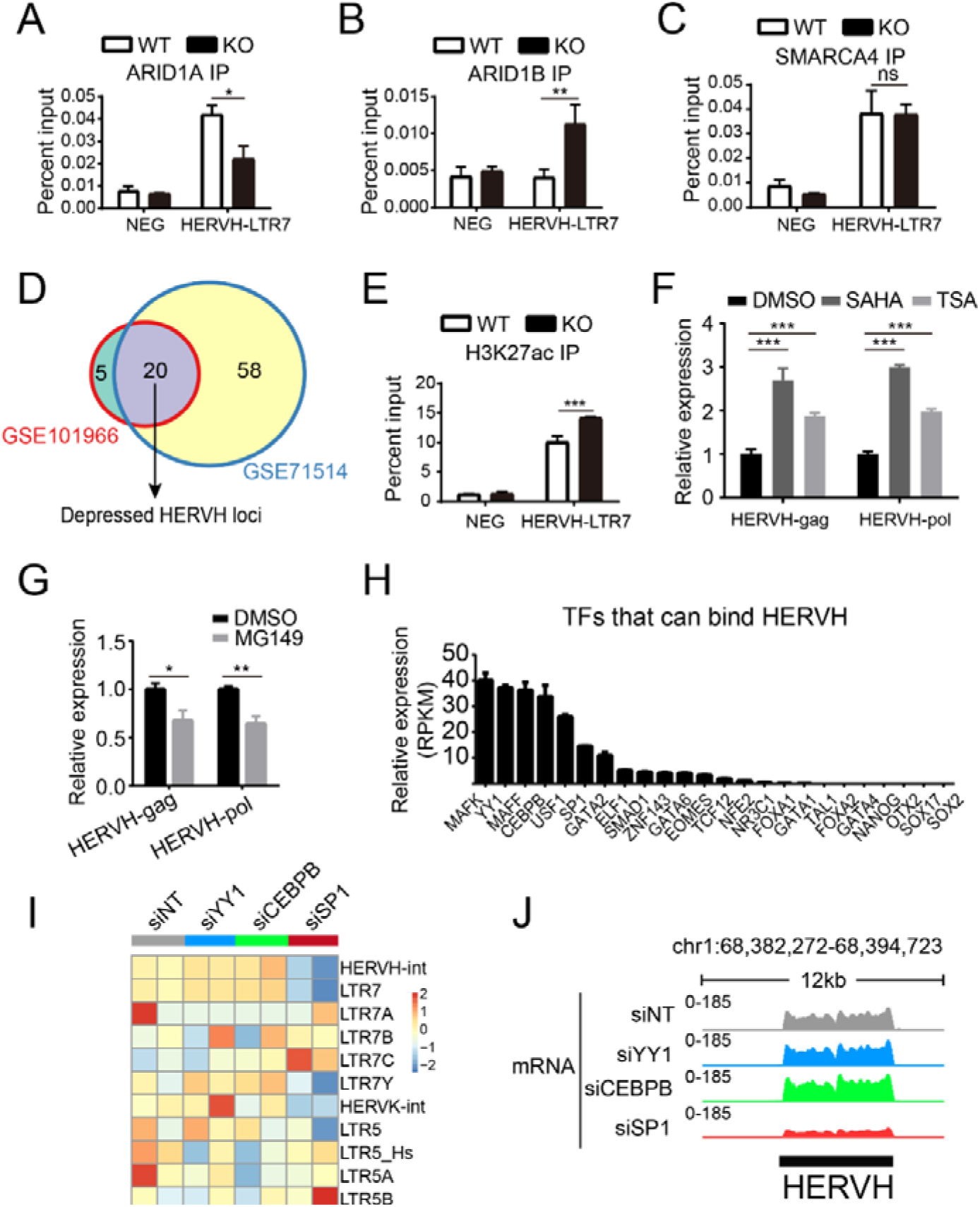
HATs associated with ARID1B and transcriptional factor SP1 are contributing to the upregulation of HERVH upon ARID1A loss. (A-C) ChIP-qPCR results showing decreased ARID1A and increased ARID1B at HERVH loci in ARID1A KO cells, whereas the amount of another BAF component SMARCA4 at HERVH loci remains unchanged. (D) Venn diagram highlights the commonly derepressed HERVH loci shared by two independent RNA-seq datasets of the ARID1A KO HCT116 cells. (E) ChIP-qPCR confirming increased H3K27ac at HERVH loci upon ARID1A loss. (F) qPCR analysis of HERVH expression in control and HDAC inhibitors treated HCT116 cells. (G) qPCR analysis of HERVH expression in control and histone acetyltransferase Tip60 inhibitor treated ARID1A KO HCT116 cells. (H) Relative expression of different transcriptional factors that are predicted to bind HERVH in ARID1A KO HCT116 cells. (H) Heatmap showing the transcripts abundance of HERVH (HERVH-int and LTR7s) and HERVK (HERVK-int and LTR5s) upon knockdown of several transcriptional factors by siRNA in ARID1A KO HCT116 cells. (J) Genomic snapshot of RNA-seq signals from ARID1A KO HCT116 cells treated with control or the indicated siRNA at a representative HERVH locus. *p < 0.05, **p < 0.01, ***p < 0.001, by t-test. ns, no significance.

Based on these results, we propose that the ARID1A-contaning BAF complex normally maintains a compact chromatin configuration at HERVH loci with the help from its associated HDACs. When ARID1A is mutated, the ARID1B-containing BAF recruits HATs to the HERVH loci and increases local accessibility, and then SP1 binds to and activates its transcription (Fig. 3I).

### HERVH is required for the survival of colorectal cancer cells

Knockdown the expression of HERVH in ESCs triggers differentiation^15^. To investigate the function of HERVH transcription in CRCs, we reduced its expression in different cell lines and patient-derived organoids using shRNAs targeting different regions of HERVH-int and assessed the consequence (Fig. S4A).

We first tested the effects of HERVH knockdown with different colorectal cell lines. The cell lines analyzed all had varying degrees of HERVH expression, whereas the expression of HERVK was kept to a minimum (Fig. S4B). Cell viability assay showed that knockdown of HERVH impaired the survival of all the cell lines tested (Fig. S4C), and the HERVH knockdown also strongly inhibited colony formation of these cells in clonogenic assays (Fig. S4D). CRC cell line SW480 had weak ability in the formation of tumor spheres when cultured in ultra-low attachment plates, and ARID1A inactivation significantly enhanced this ability (Fig. 4A). Knockdown of HERVH in the ARID1A KO cells greatly reduced the formation of tumor spheres (Fig. 4A-4C). To further assess the function of HERVH in tumorigenicity, we subcutaneously seeded shRNA-infected ARID1A WT or KO HCT116 cells into nude mice. The cells with HERVH knockdown showed significant growth impairment when compared with control cells (Fig. 4D).

**Figure 4.**
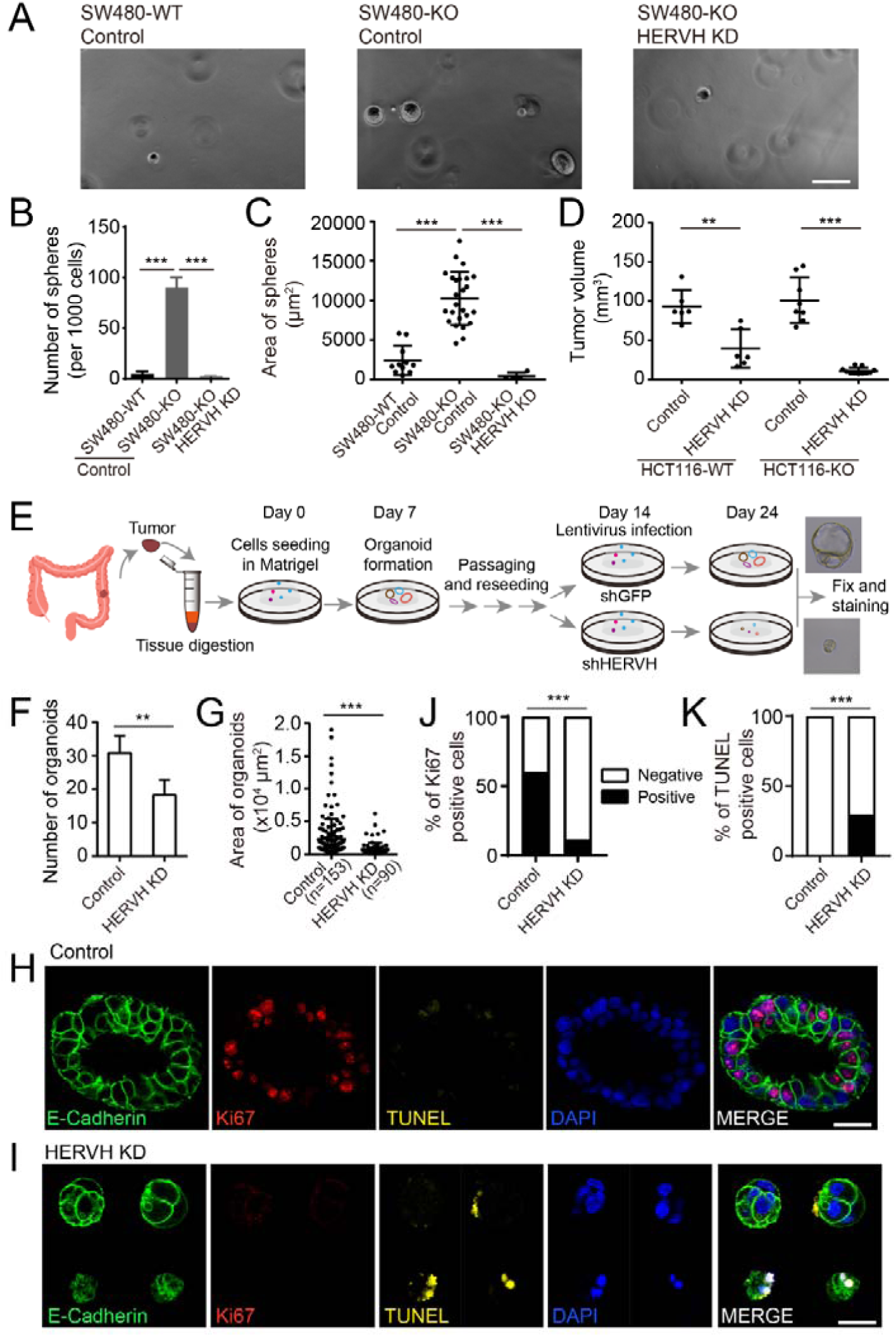
HERVH is essential for the proliferation of CRC cells. (A) Representative brightfield images showing that knockdown of HERVH inhibits sphere formation of ARID1A KO SW480 cells. Bar: 130 μm. (B-C) Quantifications of sphere number and sphere size of the indicated cells. ***p < 0.001 by t-test. (D) HERVH knockdown suppresses tumor growth of WT and ARID1A KO HCT116 cells in mouse subcutaneous xenograft tumor models. **p < 0.01, ***p < 0.001 by t-test. (E) Schematic illustrating the establishment and subsequent treatments of patient-derived CRC organoids. (F-G) HERVH knockdown decreases both the number and the size of CRC organoids. **p < 0.01, ***p < 0.001 by t-test. (H-I) Representative images of control and HERVH shRNA treated organoids stained with E-Cadherin (green), Ki67 (red), TUNEL (yellow), and DAPI (blue). Bars: 34 μm. (J-K) Percentages of Ki67 or TUNEL positive cells in control and HERVH shRNA treated CRC organoids. ***p < 0.001 by chi-squared test.

**Figure S4.**
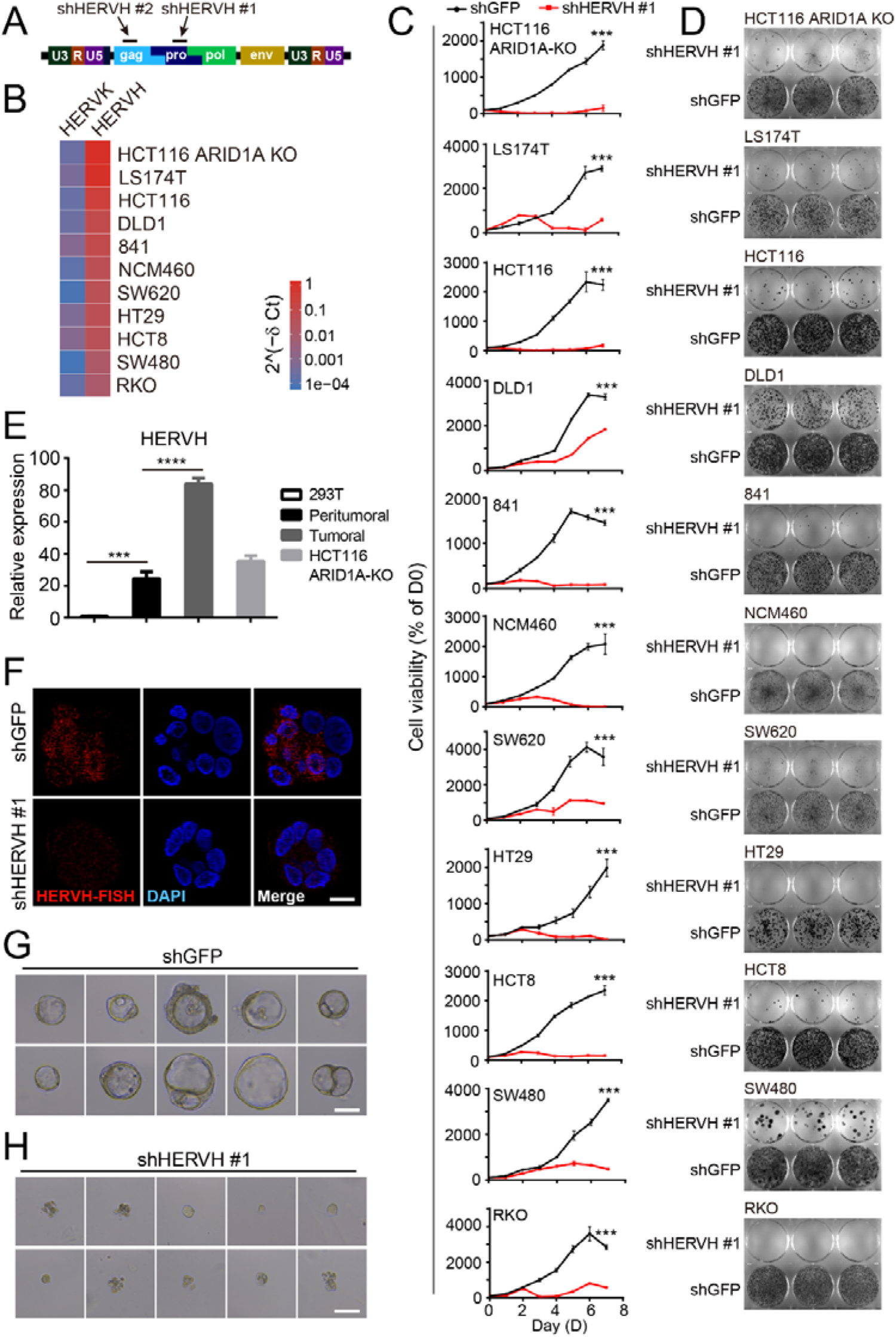
HERVH is vital for the growth of CRC cells. (A) Schematic of HERVH and the shRNAs used in this study. (B) Heatmap of qPCR results showing higher mRNA abundance of HERVH than HERVK in various normal colon and CRC cells. (C) Results of MTT assay showing reduced viability upon HERVH knockdown in the indicated cell lines. (D) Clonogenic assay showing reduced colony formation in the indicated cells treated with HERVH shRNA. (E) qPCR analysis of HERVH expression in patient samples used to establish CRC organoids. (F) Fluorescent *in situ* hybridization (FISH) shows reduced HERVH transcripts level in CRC organoids treated with HERVH shRNA. Bar: 10 μm. (G-H) Representative brightfield images of CRC organoids in control or HERVH shRNA treated groups. Bars: 74 μm. ***p < 0.001, ****p < 0.0001 by t-test.

We next sought to verify the critical role of HERVH in patient-derived CRC organoids. We obtained tumoral and peritumoral tissues from surgical biopsy, and evaluated their HERVH transcripts level using qPCR (Fig. S4E). The tumoral tissues with high HERVH expression were selected to generate CRC organoids, which were then infected with lentiviruses carrying shRNA targeting either HERVH or GFP to achieve specific knockdown (Fig. 4E). RNA fluorescence in situ hybridization (FISH) of the organoids confirmed the knockdown efficacy of HERVH (Fig. S4F). Compared to control, HERVH knockdown resulted in the formation of fewer organoids (Fig. 4F), and their size was also much smaller (Fig. S4G, S4H, and 4G). We examined the cell proliferation and apoptosis in the treated organoids by Ki67 and TUNEL stainings (Fig. 4H and 4I). HERVH knockdown dramatically reduced the number of proliferating Ki67 positive cells (Fig. 4J), meanwhile increased the number of TUNEL positive apoptotic cells (Fig. 4K).

Altogether, the results suggested that HERVH is a vulnerability not only in ARID1A mutated cells, but also in many other CRC cells that express HERVH.

### HERVH transcript is a component of BRD4 nuclear speckles and regulates BRD4-mediated transcriptions

HERVH is part of the transcriptional circuitry regulating pluripotency, and its transcription markedly influences the transcriptome^13, 15, 16, 56^. To investigate the molecular underpinnings of the oncogenic function of HERVH in CRCs, we assessed the impact of HERVH knockdown on global gene expression. PCA showed that the transcriptomes of the ARID1A WT and KO HCT116 cells were noticeably separated on the second principal component (PC2), and HERVH knockdown narrowed this difference (Fig. 5A), suggesting that the altered transcription seen in ARID1A KO cells was partially linked to the upregulation of HERVH. We identified 552 upregulated genes and 531 downregulated genes whose transcriptional change was reversed upon HERVH knockdown in ARID1A KO cells (Fig. 5B and Supplementary Table 7). Many of the 552 HERVH-dependent upregulated genes were enriched in cancer related pathways (Fig. 5C and Supplementary Table 7). We selected some of the upregulated target genes and validated the observed reversion of expression by qPCR (Fig. S5A-S5B). The observed increase of H3K27ac and H3K4me at the derepressed HERVH loci suggested that they could function as active enhancers and their transcripts enhancer RNAs^57^ (Fig. 3F and 3G). Additionally, HERVH transcripts are able to interact with many subunits of the mediator complex^15^. We compared the transcriptome dynamics after siRNA mediated knockdown of different subunits of the mediator complex, its binding partner BRD4, and HERVH. The correlation matrix suggested that suppression of BRD4 and HERVH imposed similar influence on global transcription (Fig. S5C and Supplementary Table 7). Further analysis using the differentially expressed genes in BRD4 and HERVH knockdown cells revealed strong correlation between these two groups (Fig. 5D, S5D, and Supplementary Table 7). Of the 1643 differentially expressed genes upon HERVH knockdown, 1018 of them showed similar changes in BRD4 knockdown cells (Fig. 5E and Supplementary Table 7).

**Figure 5.**
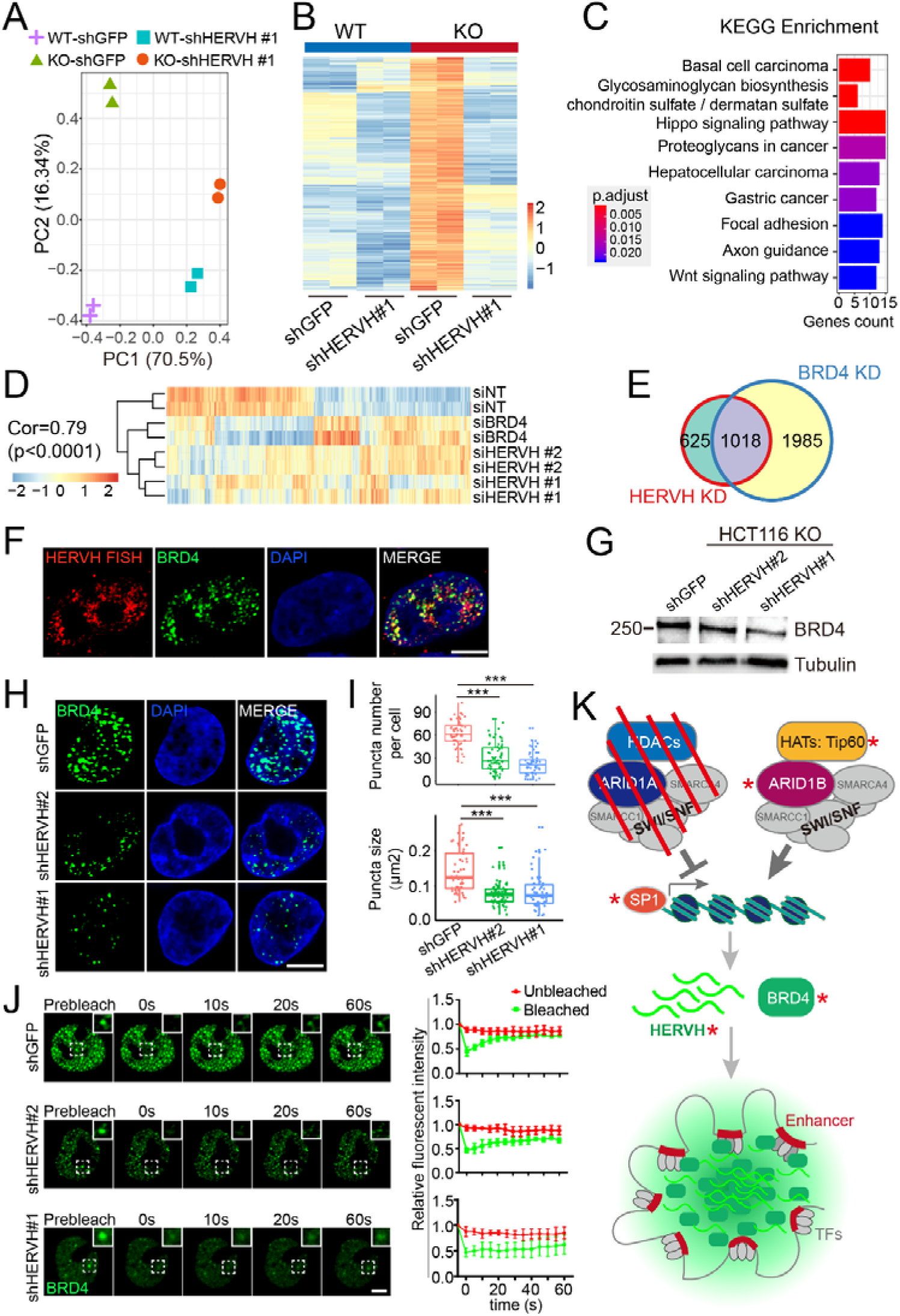
HERVH contributes to the formation of BRD4 puncta and its function in transcriptional regulation. (A) PCA analysis showing the impacts of HERVH knockdown on the global transcriptome of HCT116 WT and KO cells. (B) Heatmap highlighting a set of genes whose expression increase upon ARID1A loss but decrease again when HERVH is knocked down. (C) KEGG enrichment analysis of the HERVH-dependent genes. (D) Heatmap showing strong correlation of changes in transcriptome between knockdowns of HERVH and BRD4 in ARID1A KO HCT116 cells. (E) Venn diagram showing that 1018 genes are coregulated by BRD4 and HERVH. (F) Partial colocalization between HERVH FISH signals and GFP-BRD4 nuclear foci. Bar: 5 μm. (G) Western blot showing decreased BRD4 protein level upon knockdown of HERVH in HCT116 ARID1A KO cells. (H) Representative images of BRD4 nuclear foci in control and HERVH shRNA treated cells. Bar: 5 μm. (I) Quantifications of the number and size of BRD4 foci. ***p < 0.001 by Wilcox test. (J) Fluorescence recovery after photo bleaching (FRAP) analysis of BRD4 foci after control or HERVH knockdown. Bar: 3.3 μm. (K) A model summarizing how ARID1A loss upregulates HERVH and hence stimulates BRD4 nuclear foci formation and BRD4-mediated transcription. Red asterisks label potential targets of intervention.

**Figure S5.**
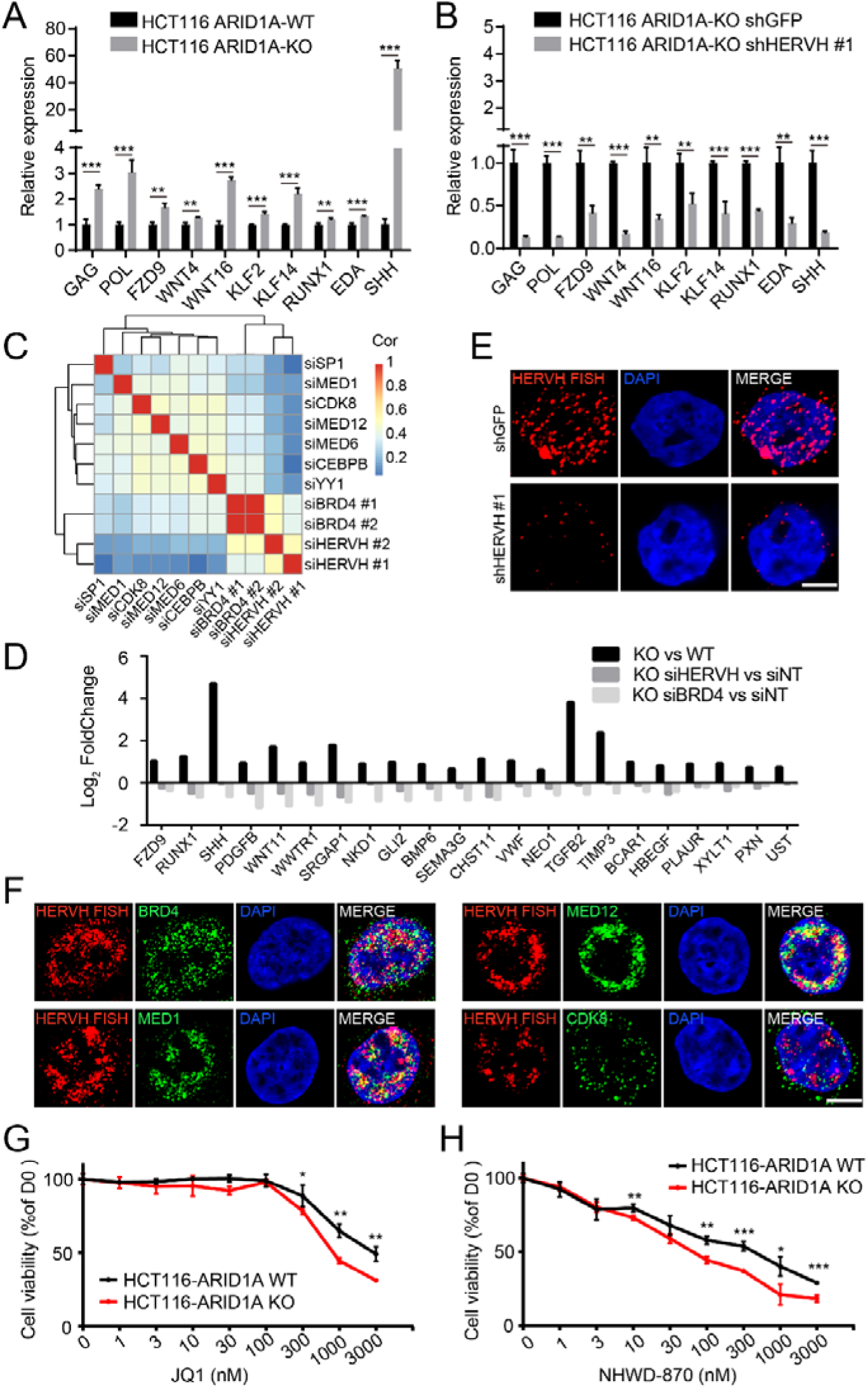
The effect of HERVH knockdown in gene expression and its relationship with BRD4 and other components of the mediator complex. (A-B) qPCR validation of gene expression changes reported in Fig 5B. (C) Correlation matrix showing the unbiased and pairwise comparisons of global transcriptome changes upon knockdown of HERVH and the indicated genes. Color bar represents Spearman’s correlation coefficient. (D) The foldchanges of a group of representative genes whose expression increases in ARID1A KO cells but decreases upon knockdown of HERVH or BRD4. (E) FISH staining of HERVH in control and HERVH knockdown cells. Bar: 5 μm. (F) Representative immunofluorescence images showing different degrees of colocalization between HERVH transcripts (red) and BRD4 or other components in the mediator complex (green). Bar: 5 μm. (G-H) MTT assay of ARID1A WT or KO HCT116 cells treated with two different BET inhibitors. *p < 0.05, **p < 0.01, and ***p < 0.001 by t test.

BRD4 as well as the mediator complex subunit MED1 can form liquid-like condensates, especially at super-enhancers^58, 59^. To investigate the distribution of HERVH transcripts and their relationship with BRD4 and the mediator complex, we combined RNA FISH targeting HERVH with immunofluorescence (IF) stainings. The specificity of the RNA FISH was validated by the absence of signals in HERVH knockdown cells (Fig. S5E). Varying degrees of colocalization was detected between HERVH transcripts and the endogenous BRD4, MED1, and MED12 (Fig. S5F), suggesting that HERVH RNA could regulate their protein dynamics in the nucleus. We knocked down the expression of HERVH, and observed a mild decrease in the protein level of BRD4 (Fig. 5G). To further characterize the influence of HERVH on BRD4, we stably expressed GFP-BRD4 in ARID1A KO HCT116 cells. GFP-BRD4 formed nuclear condensates as previously reported^58^, and RNA FISH revealed clear distribution of HERVH RNA in the BRD4 puncta (Fig. 5F). Knockdown of HERVH markedly decreased both the number and size of the BRD4 puncta (Fig. 5H and 5I). We further assessed the dynamics of the BRD4 puncta using fluorescence recovery after photobleaching (FRAP). While in control cells the photobleached BRD4 puncta quickly recovered its fluorescence, the fluorescence recovery after HERVH knockdown became much slower (Fig. 5J).

It was reported that ARID1A mutant cells showed higher sensitivity to the BET inhibitor JQ1^45, 46^. We confirmed that the ARID1A KO HCT116 cells were indeed more sensitive to JQ1 as well as another recently reported BRD4 inhibitor NHWD-870^60^ (Fig. S5G and S5H). The results reported here suggested that the upregulated transcription of HERVH in ARID1A mutant cells contributed to the formation of BRD4 nuclear puncta and stimulated their dynamic activity (Fig. 5K), providing an explanation for the observed increased sensitivity to BET inhibitors.

## Discussion

Mutational landscape analyses have revealed that ARID1A is among the most frequently mutated epigenetic factors across many cancer types^40, 44^. Understanding its mechanism of action and hence identifying targetable vulnerabilities for ARID1A inactivation have been of great importance. In this study, we investigated how the repetitive genome responded to the inactivation of ARID1A and identified that the HERVH group of ERVs was specifically derepressed. This derepression was ARID1B-dependent, and was indispensable for the survival of the CRC cells, likely due to its influence on the dynamics of BRD4 and the regulated transcriptional network. Several synthetic lethality targets of ARID1A have been reported, including ARID1B, EZH2, HDAC6, Aurora A, and GCLC^50, 54, 61–66^. ARID1A mutant cells are also hypersensitive to BET inhibitors^45, 46^, a promising class of anticancer drugs. Our results suggest that the activation of pluripotency-related HERVH is a shared mechanistic foundation of the previously observed ARID1B- and BET-vulnerabilities of the ARID1A mutated tumors. The ARID1B-HERVH-BRD4 regulatory axis and the adjunct mechanism reported here also offer several new potential targets of intervention (Fig. 5K), among which the HERVH itself is of most interest, because of its specific expression in early embryos and general silencing in most adult tissues. It is worth noting that ARID1A has been implicated in several other biological processes, some of which also involves HERVH, such as high-order spatial chromosome partitioning and tissue regeneration^15, 16, 22, 67–69^. The molecular mechanism reported here may have certain explanatory power in those scenarios as well.

Derived from ancient retroviral infections, ERVs are domesticated viral fossils in our genome whose activity is under close surveillance^1, 2, 4, 28^. Comprehensive interrogations in mouse ESCs have revealed that overlapping epigenetic pathways linked to heterochromatin formation are enlisted to suppress the transcription of ERVs. This includes DNA methylation (5-methylcytosine, 5mC), various histone modifications (H3K9me3, H3K27me3, H4K20me3, H4R3me2, and H2AK119ub), and their corresponding writers and readers^31, 32^. Reminiscent of the diversity of the process of heterochromatin formation in early embryos^70^, different families of ERVs rely on distinct epigenetic means to achieve silencing. The specific recognition of different ERVs by the various epigenetic mechanisms is in part mediated by the KRAB domain-containing zinc finger proteins (KZFPs), which can bind to specific DNA sequences in individual ERV and recruit KAP1 and other epigenetic modifiers. RNA mediated targeting mechanisms also contribute to the specific silencing of ERVs. piRNAs as well as other small RNA species are able to bring histone modifying activities to their complementary ERV loci^4, 28, 71, 72^. Our study reveals another mode of ERVs suppression which involves the SWI/SNF chromatin remodelers, further increasing the complexity of the epigenetic regulatory network constraining the expression of ERVs. The targeting mechanism for the BAF complex in silencing HERVH is currently unknown. It will be interesting to investigate the potential interactions between BAF and KZFPs or the small RNA machineries.

Accumulating evidence reveals that ERVs are co-opted to perform a wide range of biological functions. In early embryos and ESCs, ERVs serve as regulatory elements and alternative promoters to rewire the transcription network of pluripotency^13, 21^. Moreover, certain groups of ERVs become transcriptionally activated in an orderly fashion during embryogenesis^33^, functioning as enhancer or long noncoding RNAs^14, 15^, and sometimes synthesizing reverse transcriptase activity and even forming viral-like particles^73^. ERVs are also involved in many human diseases such as various types of cancer. The abnormally activated ERVs can produce long noncoding RNAs or functional polypeptides^37, 74–77^, enabling cancer cells to exploit and repurpose developmental pathways to promote malignancy^38^. Of particular note, the reactivated ERVs in cancer are extensively recruited as promoters to drive expression of many oncogenes in a process termed onco-exaptation^78, 79^. Our results reciprocally demonstrate that mutations of tumor suppressor can activate functionally important ERVs, suggesting the existence of positive feedback loops between ERVs and cancer driver genes. Future studies shall extend the analysis to other cancer driver genes and characterize these positive feedback loops more comprehensively. The establishment of a mutually reinforcing relationship between cancer driver genes and ERVs will deepen our understandings on the etiology of malignancy and throw new light on cancer treatments.

## Methods

### Data download

The TCGA dataset used in this study, including the RNA-seq BAM files, the gene raw count data (htseq-count files), and the annotated somatic simple nucleotide variation files (MuTect2 VCF) of patients with colon adenocarcinoma (COAD) and rectum adenocarcinoma (READ), were accessed through dbGaP accession number phs000178.v11.p8^48^ and downloaded using the gdc-client v1.6.0. The cinical overall survival (OS) information was obtained from Liu et al.^80^. The RNA-seq fastq files of normal and tumor tissues from another 18 CRC patients were downloaded from https://www.ncbi.nlm.nih.gov/geo under the accession number GSE50760^81^. The RNA-seq fastq files of the 59 colorectal cancer cell lines in cancer cell line encyclopedia (CCLE) were downloaded from https://www.ebi.ac.uk/ under the accession number PRJNA523380^49^, and the corresponding germline filtered CCLE merged mutation calls were acquired from https://portals.broadinstitute.org/ccle/data. The previously published RNA-seq and ChIP-seq raw reads fastq files generated with HCT116 cells or mice primary colon epithelial cells were downloaded from https://www.ncbi.nlm.nih.gov/geo under the accession numbers GSE71514 and GSE101966^42, 50^.

### RNA-seq analysis

Raw reads were first cleaned using trim_galore v0.6.0 (http://www.bioinformatics.babraham.ac.uk/projects/trim_galore/) with default parameters. The reads from each RNA-seq sample were then mapped to hg38 or mm9 genome assembly downloaded from UCSC, using STAR v2.5.3a^82^. The key alignment parameters were as follows: “--outFilterMismatchNoverLmax 0.04 --outSAMtype BAM SortedByCoordinate --outFilterMultimapNmax 500 --outMultimapperOrder Random --outSAMmultNmax 1”; the parameters “--outFilterMultimapNmax 500” and “--outMultimappedOrder Random” ensured that multiple aligned reads were included but only one position was assigned randomly. Genes expression was quantified using featureCounts v1.6.5^83^ of subread-1.6.5 package based on hg38 RefSeq genes annotation file. Repeats expression was quantified using featureCounts v1.6.5 (“featureCounts --M --fraction”) based on repeats annotation files downloaded from https://genome.ucsc.edu/cgi-bin/hgTables. Principal component analysis was conducted with the functions “vst” and “plotPCA” from R package DESeq2 v1.22.2^84^. Differential expression analysis was performed based on the negative binomial distribution using the functions “DESeq” and “results” from DESeq2. The heatmap of differentially expressed genes or repeats was created using R package pheatmap v1.0.12. The KEGG enrichment analysis was performed using the function “enrichKEGG” from the R package clusterProfiler v3.10.1^85^. Venn diagrams were prepared with the R package Vennerable and venn.

### Survival analysis

The curated clinical endpoint results (OS event and OS event times) of the 628 patients in TCGA-COREAD dataset were obtained from Liu et al.^80^. Only patients in stages II and later according to the American Joint Committee on Cancer (AJCC) pathologic tumor staging system were included. The 493 CRC patients were classified into HERVH-high (145 patients with HERVH-int CPM>8430.797) and HERVH-low groups (348 patients with CPM<8430.797), and the survival curves of the two groups were compared using log-rank test from the function “survdiff” in R package survival v2.44-1.1.

### Integration analysis of whole-exome sequencing (WXS) and RNA-seq

WXS files (MuTect2 VCF) and RNA-seq data from 516 patients in TCGA-COREAD were analyzed (Fig. S1A). All the somatic mutational information was included regardless of their classification. For each gene, we classified the patients into WT or mutation group, and then calculated the Log_2_ FoldChange between these two groups using the expression values (CPM) of HERVK-int and HERVH-int. p values were calculated by Wilcoxon test.

### ATAC-seq and ChIP-seq analyses

Raw reads were cleaned using trim_galore. The reads were then aligned to the hg38 genome assembly using Bowtie2 v2.3.5.1^86^, with the default parameters that look for multiple alignments but only report the one with best mapping quality. Duplicate reads were then removed using MarkDuplicates from gatk package v.4.1.4.1. Replicate samples were merged using the samtools v1.10^87^. Peak calling was performed using MACS2 v2.2.6^88^ (parameters: -g hs --keep-dup 1 --broad-cutoff 0.01). Peaks near active HERVH loci were identified using bedtools v2.26.0^89^. For ATAC-seq, bigwig tracks were generated using bamCoverage from python package deeptools (parameteres: --skipNAs --normalizeUsing CPM)^90^. For ChIP-seq, bigwig tracks were generate using bamCompare from deeptools (parameters: --skipNAs --scaleFactorsMethod readCount --operation log2 --extendReads 200). Negative values were set to zero. ATAC-seq and ChIP-seq profiles were created by computeMatrix and plotProfile in deeptools. IGV v.2.4.13 was used to visualize the bigwig tracks^91^.

### Cell culture and cell line generation

The cell lines used in this study, including HCT116, DLD1, SW480, LS174T, SW620, HT29, HCT8, RKO, CRL1790/841, NCM460, and 293T, were cultured in RPMI 1640 or DMEM medium containing 10% FBS and incubated at 37 °C with 5% CO_2_ in a humidified incubator. To generate ARID1A KO cell lines, the indicated cells were transfected with LentiCRISPR-V2 plasmid carrying sgARID1A (Supplementary Table 8) using Lipofectamine 2000 (Invitrogen), and further selected by 1 μg/mL puromycin (Selleck, s7417) for 3 days. The cells were then plated at single-cell density in 100 mm petri dishes, and the emerged individual clones were picked and replated into 24-well plates. The loss of ARID1A expression was confirmed by western blot.

### Organoid culture

The CRC organoid was generated as previously described^92^. All the human tissue related experiments were approved by the Medical Ethics Committee of Central South University, and the informed consent was obtained from the patients. From the resected colon segment, the tumor tissues as well as normal tissues were isolated and stored in ice-cold RPMI 1640 supplemented with 1% Penicillin-Streptomycin. The tissues were then washed in ice-cold DPBS supplemented with 1% Penicillin-Streptomycin and cut into 1-3 mm^3^ cubes. After centrifuging at 200 g for 5 min, the supernatant was removed and pellet was resuspended in collagenase IV (Gibco, 17104019) supplemented with 10 µM ROCK inhibitor Y-27632 dihydrochloride (Merk Millipore, SCM075). The tissues were digested at 37□ for 1 hour and mixed up every 10-15 min by pipetting, washed with 10 mL advanced DMEM/F12 (Thermo Fisher Scientific, 12634-010) supplemented with Y-27632, and then centrifuged at 200 g for 5 min at 4 °C. The pellet was resuspended in DMEM/F12 supplemented with Y-27632 and filtered through 60 µm cell strainer. After centrifugation at 200 g for 5 min at 4 °C, the supernatant was discarded and the pellet resuspended in 70% Matrigel (Corning, 356231). 30 µL of the Matrigel mixture was plated on the bottom of 24-well plates, and 500 µL organoid medium (Accurate International Biotechnology, M102-50) was added to each well following incubation at 37□ with 5% CO_2_ for 30 min. The organoid medium was changed every 2-3 days, and the organoids were passaged after 7 days of culture.

### Cell growth assays

For cell viability assays, cells were plated into 96-well plates at the density of 2000-5000 cells per well after infected with lentiviruses expressing shGFP or shERVH. The cells were kept for another 7 days, and the viability was measured daily using MTT (Sigma, M5655) as previously described^93^. For chemosensitivity assays, the cells were seeded in 96-well plates and treated with the compounds at indicated concentrations for 72 hours, and then the cell viability was measured. For colony formation assays, the cells were seeded at the density of 1000-2000 cells per well in 6-well plates after infected with lentiviruses expressing shGFP or shERVH. The cells were allowed to grow for 10-14 days and then fixed for 10 min in 50% (v/v) methanol containing 0.01% (w/v) crystal violet.

### Tumor sphere formation

The 6-well plates were coated with 12 mg/mL poly-hydroxyethylmethacrylate (polyHEMA, Sigma-Aldrich, P3932) in 95% ethanol. The indicated cells were digested by TrypLE, and approximately 1000 cells were suspended in 50% Matrigel (Corning, 356231) and plated in the precoated 6-well plates. The 6-well plates containing the cells were incubated at 37□ for 30 min, and then 2 mL of phenol red-free DMEM/F12 (GIBCO, 21041) containing 1× B27 supplement (Invitrogen, 12587) and 20 ng/mL rEGF (Sigma Aldrich, E-9644) was added into each well. The culture medium was changed every 2-3 days, and the number of tumor spheres in each well was counted after 12 days.

### Xenograft tumors

The 4-5 weeks old female BALB/c nude mice were purchased from Hunan SJA Laboratory Animal Co., Ltd. (Changsha, China). 5×10^5^ of the indicated cells were suspended in 100 µL DPBS and injected subcutaneously into the flank of nude mice. The tumors were measured twice weekly with an electronic caliper, and the volumes were calculated using the formula: 0.5×(length × width^2^). All the animal experiments were approved by the Medical Ethics Committee of Central South University, and conducted according to the Guidelines of Animal Handling and Care in Medical Research in Hunan Province, China.

### RNA interference

The siRNA oligos were synthesized by GenePharma (Shanghai GenePharma Co., Ltd.), and the sequences were listed in Supplementary Table 8. Cells were transfected with the indicated siRNA by Lipofectamine 2000. After 48 hours, the cells were harvested and the efficiency of silencing was verified by qPCR. For shRNA, shRNA oligos were synthesized by Tsingke (Tsingke Biotechnology Co., Ltd.) and cloned into pLKO.1 TRC Cloning vector (Supplementary Table 8). The shRNA and packaging vectors (pMD2.G and psPAX2) were transiently co-transfected into 293T cells by polyethylenimine (Sigma, P3143), and the resulted lentivirus particles were harvested and precipitated by PEG8000. The target cells were treated with lentivirus particles and 8 µg/mL polybrene for 24 hours, and the efficacy of shRNA interference was determined by qPCR.

### HERVH knockdown in organoids

The organoids cultured in Matrigel were washed once with DPBS, and digested with TrypLE for 5 min at 37□. During the digestion, Matrigel was disrupted by pipetting repeatedly. When cell clumps containing 2-10 cells were observed, 10 mL of advanced DMEM/F12 was added before centrifugation at 200 g for 5 min. The supernatant was removed and the cells were resuspended using organoid medium supplemented with 8 µg/mL polybrene. Then the cells were split equally into 2 wells of 24-well plate precoated with polyHEMA, and 50 µL of lentivirus carrying shGFP or shERVH was added. After spin infection at 2000 rpm for 1 hour, the cells were incubated at 37□ with 5% CO_2_ for 4 hours. The cells were then resuspended in 10 mL of advanced DMEM/F12 and centrifuged at 200 g for 5 min. The pellet was resuspended with 100 µL of 70% Matrigel, and 10 µL of the mixture was plated per well into prewarmed 96-well plate. The organoids were cultured for 10-14 days and the medium was changed every 2-3 days.

### Western blot

Cells were washed with cold DPBS for two times and then lysed in 2× Laemmli buffer (2% SDS, 20% glycerol, and 125 mM Tris-HCl, pH 6.8) supplemented with 1× protease inhibitor cocktail (Sigma, P8340). The cell lysate was scraped and sonicated, and the concentration of protein was determined by BCA assay. The protein was separated by SDS-PAGE and transferred onto nitrocellulose membrane. The membrane was then blocked with 5% non-fat milk for 1 hour at room temperature, and incubated with the indicated primary antibody overnight at 4□ with shaking. The membrane was washed for 3 times and incubated with secondary antibodies (1:5000, Thermo Fisher Scientific) for 2 hours. The signal was then detected with ECL substrates (Millipore, WBKLS0500). Dilutions of primary antibodies were: rabbit anti-ARID1A/BAF250A Ab (1:1000, Cell Signaling, 12354S), rabbit anti-BRD4 Ab (1:1000, Active Motif, 39909), mouse anti-α-Tubulin Ab (1:3000, Cell Signaling, 3873s). Primary antibodies used in this study were listed in Supplementary Table 11.

### RNA-seq and qPCR

The RNA of the treated cells was extracted by TRIzol (Life Technologies, 87804) according to the manufacturer’s protocol. Total RNA was made into libraries for sequencing using the mRNA-Seq Sample Preparation Kit (Illumina) and sequenced on an Illumina Hiseq platform (Novagene, Tianjin, China). The sequencing data was deposited to the GEO database (accession number GSE). For RT-qPCR, RNA was extracted by TRIzol, and reverse transcribed to cDNA using the PrimeScript RT reagent Kit (Takara, RR037A). The cDNA was then used as templates and qPCR was performed using the SYBR Green qPCR Master Mix (SolomonBio, QST-100) on the QuantStudio 3 Real-Time PCR system (Applied Biosystems). Primers used in qPCR were listed in Supplementary Table 8.

### Chromatin immunoprecipitation

The indicated cells in 100 mm petri dishes were cross-linked with 1% formaldehyde for 10 min at room temperature, and quenched with 125 mM ice-cold glycine. The cells were then rinsed with 5 mL ice-cold 1× PBS for two times, and harvested by scraping using silicon scraper. After spinning at 1350 g for 5 min at 4□, the supernatant was discarded, and the pellet was resuspended in Lysis Buffer I (50 mM HEPES-KOH, pH 7.5, 140 mM NaCl, 1 mM EDTA, 10% glycerol, 0.5% NP-40, 0.25% Triton X-100 and 1× protease inhibitors) and incubated at 4□ for 10 min with rotating. After spinning at 1350 g for 5 min at 4□, the pellet was resuspended in Lysis Buffer II (10 mM Tris-HCl, pH 8.0, 200 mM NaCl, 1 mM EDTA, 0.5 mM EGTA and 1× protease inhibitors), incubated for 10 min at room temperature, and spun at 1350 g for 5 min at 4□. The pellet was again resuspended in Lysis Buffer III (10 mM Tris-HCl pH 8.0, 100 mM NaCl, 1 mM EDTA, 0.5 mM EGTA, 0.1% Na-Deoxycholate, 0.5% N-lauroylsarcosine and 1× protease inhibitors) and transferred into Covaris microTUBEs. The DNA was sonicated to 200 bp fragments using Covaris S220 (duty cycle: 10; intensity: 4; cycles/burst: 200; duration: 200 s). After quenching the SDS by 1% of Triton X-100, the lysate was spun at 20,000 g for 10 min at 4□. 50 µL of supernatant from each sample was reserved as input, and the rest lysate was incubated overnight at 4□ with the magnetic beads bound with ARID1A (CST, 12354S), ARID1B (Santa Cruz, sc-32762X), SMARCA4 (Abcam, ab110641) or H3K27ac (Abcam, ab4729) antibody respectively. The beads were washed three times with Wash Buffer (50 mM Hepes-KOH, pH 7.6, 500 mM LiCl, 1 mM EDTA, 1% NP-40, 0.7% Na-deoxycholate), and washed once with 1 mL TE buffer containing 50 mM NaCl. The DNA was eluted with 210 µL of Elution Buffer (50 mM Tris-HCl, pH 8.0, 10 mM EDTA, 1% SDS). The cross-links were reversed by incubated at 65□ overnight. 200 µL of TE buffer was added to each tube, and the RNA was degraded by incubation with 16 µL of 25 mg/mL RNase A at 37□ for 60 min. The protein was degraded by adding 4 µL of 20 mg/mL proteinase K and incubating at 55 °C for 60 min. The DNA was then purified by phenol:chloroform:isoamyl alcohol extraction, and resuspended in 50 µL ddH_2_O. The fragments of HERVH DNA were detected by qPCR (Supplementary Table 8).

### The RNAscope™ in situ hybridization (ISH)

The colon cancer tissue array (HCol-Ade180Sur) was purchased from Shanghai Biochip Co. Ltd (Shanghai, China). The RNAscope analysis with probes targeting the HERVH-gag sequence was performed using the RNAscope Multiplex Fluorescent Reagent Kit v2 (ACD bio, 323100) according to the manufacturer’s protocol. The HERVH consensus sequence used for probe design was listed in Supplementary Table 9. Following the RNAscope staining, the tissue array was imaged with a LSM880 confocal microscope (Zeiss).

### RNA-FISH combined with immunofluorescence

RNA-FISH combined with immunofluorescence was performed as previously described^58^. Cells cultured on poly-L-lysine-coated coverglasses were fixed with 10% formaldehyde in DPBS for 10 min. After three washes in DPBS, cells were permeabilized with 0.5% Triton-X100 for 10 min. The cells were then washed three times in DPBS and blocked with 4% Bovine Serum Albumin for 30 min. The cells were incubated with the indicated primary antibody diluted in DPBS overnight, washed three times in DPBS, and incubated again with the secondary antibody for 1 hour. After washing twice with DPBS, the cells were fixed again with 10% formaldehyde in DPBS for 10 min. Following two washes with DPBS, the cells were further washed in Wash Buffer I (20% Stellaris RNA FISH Wash Buffer A (Biosearch Technologies, Inc., SMF-WA1-60), 10% Deionized Formamide (Invitrogen, AM9342) in RNase-free water) for 5 min. The RNA probe (Stellaris) in hybridization buffer was added to the cells and incubated at 37□ for 16 hours. After washing with Wash Buffer I at 37□ for 30 min, the cells were stained with 1 µg/mL DAPI for 5 min. The cells were then washed with Wash Buffer B (Biosearch Technologies, Inc., SMF-WA1-60) for 5 min, and rinsed once in water before mounting with SlowFade Diamond Antifade Mountant (Invitrogen, S36963). The sequence of the RNA probe (Stellaris) was listed in Supplementary Table 10.

### RNA-FISH and immunofluorescence with organoids

After dissolving the Matrigel with ice-cold cell recovery solution (Corning, 354253), the organoids were placed on a poly-L-lysine-coated glass slide for 30 min. The organoids attached to the slide were fixed with 10% formaldehyde for 45 min at 4 °C, and washed with DPBS for three times. The organoids were then permeabilized with 0.5% Triton-X100 for 15 min and washed with DPBS for two times. After one wash with Wash Buffer A for 5 min, the organoids were hybridized with the RNA-FISH probe overnight at 37□. After one wash with Wash Buffer A for 30 min at 37□, the organoids were stained with 1 µg/mL DAPI in Wash Buffer A for another 30 min, and washed twice with Wash Buffer B for 30 min. The organoids were rinsed with ddH_2_O and mounted with SlowFade Diamond Antifade Mountant (Invitrogen, S36963). The images were taken with a LSM880 confocal microscope (Zeiss).

The immunofluorescence of organoids was performed as previously described^94^. The organoids cultured in 96-well plate were washed once with DPBS without disrupting the Matrigel, and then 200 µL of ice-cold cell recovery solution (Corning, 354253) was added and incubated at 4 °C for 1 hour with shaking at 60 rpm. After the Matrigel was dissolved, the organoids were rinsed out using ice-cold PBS with 1% BSA and spun down at 70 g for 3 min at 4 °C. The pellet of organoids was resuspended in 1 mL of 10% formaldehyde in DPBS, and incubated at 4 °C for 45 min. 9 mL of ice-cold PBT (0.1% Tween 20 in DPBS) was added and incubated at 4 °C for 10 min. The organoids were then spun down at 70 g for 5 min at 4 °C, resuspended in 200 µL ice-cold OWB (0.1% Triton X-100, 0.2% BSA in DPBS), and transferred into 24-well plate precoated with polyHEMA. Following incubation at 4 °C for 15 min, 200 µL of the indicated primary antibody diluted in OWB was added and incubated overnight at 4 °C with shaking at 60 rpm. The next day, 1 mL of OWB was added into each well. After all the organoids were settled at the bottom of the well, the OWB was removed with just 200 µL left in each well. The organoids were washed three times with 1 mL of OWB and incubated at 4 °C for 2 hours with shaking at 60 rpm. The OWB was removed with just 200 µL left in each well, and then 200 µL of secondary antibody diluted at 1:200 in OWB was added and incubated overnight at 4 °C with shaking at 60 rpm. After the incubation, the organoids were washed once with OWB, and 200 µL of 2 µg/mL DAPI in OWB was added and incubated at 4 °C for 30 min. The organoids were then washed two times with OWB, transferred to 1.5-mL Eppendorf tube, and spun down at 70 g for 3 min at 4 °C. The OWB was removed as much as possible without touching the organoids, and the organoids were resuspended with fructose-glycerol clearing solution (60% glycerol and 2.5 M fructose in ddH_2_O). Drew a 1×2 cm rectangle in the middle of a slide, and placed 3 layers of sticky tape at both sides of the rectangle. The organoids were transferred into the middle of the rectangle, and put the coverslip on the top. The images were taken with a LSM880 confocal microscope (Zeiss).

### Fluorescence Recovery After Photobleaching (FRAP)

The treated cells were plated into 35 mm glass bottom confocal dishes (NEST, 801001), and the FRAP experiment was performed on the Zeiss LSM880 Airyscan confocal microscope with a 63x Plan-Apochromat 1.4 NA oil objective. The Zeiss TempModule system was used to control the temperature (37 °C), the humidity and the CO_2_ (5%) of the imaging system. After imaging for 3 frames, the cells were photo-bleached using 100% laser power with the 488 nm laser (iterations: 50, stop when intensity drops to 50%). The cells were then imaged again every two seconds. The images were analyzed and measured with ZEN 2 blue edition (Zeiss).

### Code Availability

All custom scripts are available from the authors upon request.

## Acknowledgements

We gratefully acknowledge Prof. François Mallet for HERVH antibodies, and Prof. Xiang Chen and Prof. Mingzhu Yin for BET inhibitors. We thank Dr. Kai Fu and Dr. Joong Sup Shim for cell lines, and colleagues in the center of medical genetics in Central South University and members of the Yuan lab for helpful discussion. This project has been supported by Central South University (2018CX032 to K.Y, 2019zzts339 to X.L, and the innovation-driven team project 2020CX016), Department of Science & Technology of Hunan Province (grants 2017RS3013, 2017XK2011, 2018DK2015, and 2019SK1012 to K.Y, 2019JJ40478 to P.L, and the innovative team program 2019RS1010), the National Natural Science Foundation of China (grants 31771589 and 91853108 to K.Y, 81801426 to L.S). K.Y. is supported by the National Thousand Talents Program for Young Outstanding Scientists.

## Author contributions

C.Y., X.L., X.H., P.L., and K.Y. conceived the project. X.H. initiated this project. C.Y., X.L., F.C., S.M., L.L., H.L., X.H., R.W., L.S., N.Z., Y.M., Y.S., and P.L. performed the experiments. X.L. and S.M. performed all the bioinformatic and statistical analyses with the supervisions from K.Y. and P.L. C.L. assisted with patient selection. S.H., H.S., and Z.Z. provided insightful suggestions in project design and manuscript writing. K.Y. wrote the manuscript with helps from C.Y. and X.L.

